# Shifted dynamics of glucose metabolism in the hippocampus during aging

**DOI:** 10.1101/2021.04.25.441340

**Authors:** Ivan Ge, Gregory W. Kirschen, Xinxing Wang

## Abstract

Aging is a process that adversely affects brain functions such as cognitive behaviors. Brain activity is a high-energy consumption process. Glucose serves as the main energy source under normal circumstances. Whether the dynamics of glucose metabolism during aging remains unchanged is not well understood. This study sought to investigate the activity-dependent changes in glucose metabolism of the mouse hippocampus during aging. In brief, after one hour of contextual exploration in an enriched environmental condition or one hour in a familiar home cage condition, metabolites were measured from the hippocampus of both adult and aged mice with metabolomic profiling. Compared to the home cage context, the enriched contextual exploration condition resulted in changes in the concentration of 11 glucose metabolism-related metabolites in the adult hippocampus. In contrast, glucose metabolism-related metabolite changes were more apparent in the aged group during contextual exploration when compared to those in the home cage condition. Importantly, in the aged groups, several key metabolites involved in glycolysis, the TCA cycle, and ketone body metabolism accumulated, suggesting the less efficient metabolization of glucose-based energy resources. Altogether, the analyses revealed that in the aged mice during enriched contextual exploration, the glucose resource seems to be unable to provide enough energy for hippocampal function.

**Highlights:**

1. Glycolysis becomes less efficient in the aged hippocampus during contextual exploration.
2. TCA cycle is altered in the aged hippocampus during contextual exploration.
3. Ketone body metabolism is elevated in the aged hippocampus during contextual exploration.

## Introduction

Aging is a degenerative process that affects almost all biological organisms (Mattson and Arumugam, 2018). During the aging process, there is a buildup of biological waste products and metabolites that likely contribute to the development of degenerative diseases. For instance, aging has been associated with diseases such as atherosclerosis with the accumulation of oxidized lipid particles in the subintimal layer of the arterial wall (Tyrrell and Goldstein, 2021), and age-related macular degeneration with the accumulation of cholesterol-rich drusen deposits in the central area of the retina (Al-Zamil and Yassin, 2017). Importantly, age-related degeneration has a particularly prominent effect on the brain and can negatively impact important functions such as cognition and sleep (Mattis and Sehgal, 2016; Mander et al., 2017; Schaum et al., 2020). Although many molecular mechanisms underlying brain degeneration have been discovered, the mechanisms involved in possible changes in energy-related metabolism remain largely unknown.

Since neurons require high energy consumption to function normally, notably to drive a sodium gradient across the cell membrane via the Na-K-ATPase, the cell’s energy production capability is vital for the execution of brain physiological functions (Dienel, 2019). Although previous research demonstrates that energy metabolism is altered in the aged brain compared to the adult brain, a more specific and quantitative description of energy metabolic pathways in the brain (Hoyer, 1990; Camandola and Mattson, 2017; Drulis-Fajdasz et al., 2018; Manza et al., 2020), especially in animals completing behavioral tasks, is not well understood. Glucose is the major energy resource responsible for supporting brain activity under non-starvation conditions (Mergenthaler et al., 2013; Dienel, 2019). Previous non-invasive imaging techniques provided direct and indirect evidence of glucose level changes in the brain during neural activity (Choi et al., 2001; Raichle and Mintun, 2006; Magistretti and Allaman, 2015). It is also important to note that several other studies showed that activity led to increases in blood flow and glucose, but only slightly increases oxygen consumption. This elucidates the hypothesis that both oxidative and nonoxidative glucose metabolism might be involved in providing energy for neural activity (Belanger et al., 2011; Figley and Stroman, 2011). However, detailed metabolic mechanisms underlying these changes are lacking.

To better understand the possible changes in metabolites during hippocampal activity, we used a metabolomics method to analyze the dynamics of energy-related pathways in the mouse brain. In this present study, we performed a global metabolomics analysis to investigate the differences in the hippocampal energy metabolism of both adult (6-week-old) and aged (78-week-old) under resting and hippocampal-engaged contextual exploration conditions. Metabolites in the glucose metabolic pathway, tricarboxylic acid (TCA) cycle, and ketone body metabolism were measured in both age categories as well as both conditions. The activity-dependent dynamics were compared between the two conditions. Strikingly, we found substantially more changes in concentrations of metabolites in the aged group during contextual exploration compared to that of the adult group. Interestingly, in the aged group, the glycolytic efficiency decreased while the ketone body metabolism increased. While changes in the overall energy production remain unknown, a clear shift in the energy production pathway was observed.

## Materials and Methods

### Experimental Animals

Mice used for the metabolomics were either 6-week (adult) or 78-week (aged) C57BL/6 male mice (The Jackson Laboratory). Animals were housed under a 12-hour light/dark cycle cage and given *ad libitum* access to food and water. All experimental procedures were carried out in accordance with guidelines from the National Institutes of Health and approved by the Stony Brook University Animal Care and Use Committee.

### Enriched contextual exploration

Adult and aged mice cohorts that were set as a contextual exploration group were given one week to become familiar with the exploration task. Animals were exposed in a new cage with 4-6 novel objects (EE) for an hour for each training. In contrast, adult and aged animal cohorts that were always kept in their home cages without extra objects were set as control groups (HC). During the night cycle when all mice were active, all animals were sacrificed in a random order.

### Metabolomics profiling

Six 6-week-old mice were set as an adult group and six 78-week-old mice were set as an aged group. Animals were anesthetized with urethane (i.p. 1.5g/kg) and transcardially perfused with ice-cold phosphate-buffered saline (PBS) buffer. In each mouse, two hippocampi were separated immediately from both hemispheres on ice and directly frozen by dry ice before being stored in a - 80°C freezer until processing. For sample extraction, several recovery standards were used before the first step in the extraction process to allow confirmation of extraction efficiency (MicroLab STAR system, Hamilton Company). To remove protein, small molecules bound to protein were dissociated or trapped in the precipitated protein matrix, and chemically diverse metabolites were recovered, with samples precipitated with methanol for 2 minutes (Glen Mills GenoGrinder 2000) followed by centrifugation. Samples were then placed briefly on a TurboVap (Zymark) to remove the organic solvent content and then frozen and dried with nitrogen. Instrument variability was determined by calculating the median relative standard deviation for the standards added to each sample before injection into the mass spectrometers. Overall process variability was determined by calculating the median relative standard deviation for all endogenous metabolites (i.e., non-instrument standards) present in 100% of the pooled matrix samples. Metabolomic profiling analysis utilized multiple platforms, including ultra-high performance liquid chromatography/tandem mass spectrometry (UPLC-MS/MS) methods and hydrophilic interaction chromatography (HILIC)/UPLC-MS/MS, which was performed by Metabolon, Inc. Aliquots of the sample were analyzed using a Waters Acquity UPLC (Waters Corp.) and LTQ mass spectrometer (MS) (Thermo Fisher Scientific, Inc.), which consisted of an electrospray ionization source and a linear ion-trap mass analyzer. The MS analysis alternated between MS and data-dependent MSn scans using dynamic exclusion; scanning varied slightly between methods but covered 70–1,000 mass-to-charge ratio.

A global metabolite class within carbohydrates and energy categories was investigated. Compounds were identified by comparison to library entries for purified standards, including retention time/index, mass-to-charge ratio, and chromatographic data (including MS/MS spectral data) using Metabolon’s hardware and software. Peaks were quantified using the area under the curve. Each raw concentration of metabolite was rescaled to set the median equal to 1, and missing values were imputed with the minimum. Comprehensive metabolomic data analysis was performed using MetaboAnalyst 5.0 (https://www.metaboanalyst.ca). Principal component analysis (PCA) was performed using MATLAB and applied for a preliminary evaluation of data quality between groups. Hierarchical cluster analysis (based on t-tests) was performed to create a heatmap of differentially expressed metabolites using MetaboAnalyst 5.0.

### Statistical analysis

The original values for each metabolite are normalized in terms of raw area counts. All metabolites are rescaled to set the median equal to 1 and the missing values are imputed with the minimum. The data was analyzed with Student’s t-tests (unpaired, two-tailed). A value of p < 0.05 was considered statistically significant. All data are presented as mean ± standard error of the mean (SEM).

## Results

### Metabolomics of the aged hippocampus during an enriched contextual exploration

Endogenous neurogenesis, neural circuit plasticity, and cognitive function in the hippocampus all decline during aging (Mattson and Arumugam, 2018). Recent studies, including the work from our group, have shown that the hippocampus, a key brain area for episodic memory, exhibits dramatic changes in concentration of metabolites related to energy consumption during a behavior task (McNay et al., 2000; Wang et al., 2020). While previous studies have shown the declining functionality of the aging hippocampus, the causes underlying this decay remain largely unknown. The impaired energy supply may contribute to the dysfunction of the aged hippocampus given that neuronal coding is an energy-intensive activity. We, therefore, aim to analyze the dynamics of energy production-related metabolites in the aged hippocampus in comparison to those of the adult hippocampus during enriched contextual exploration, which serves as a stimulus to activate the hippocampus.

To analyze the dynamics of these metabolites in the hippocampus across aging, we first profiled metabolites of the hippocampus during hippocampus-engaged behaviors as outlined in **Figure 1A**. We arranged a cohort of 12 male mice that were 6-weeks-old and another cohort of 12 male mice that were 78-weeks-old. Six of the mice of each age were housed in a cage with 5-6 novel objects, defined as an enriched environment group (EE), to increase context exploration as we have previously described (Shen et al., 2019; Wang et al., 2020). The other six mice of each age were placed in a cage with only bedding, defined as a home cage group (HC or resting group). One hour after the mice were left in these two conditions, we sacrificed the mice and promptly collected the entire hippocampi for metabolomics analysis. At both ages, out of the 593 analyzed metabolites, we sorted out 54 metabolites closely linked with carbohydrate metabolism and energy production. Using principal component analysis (PCA) of six biological replicates of each group, we found a clear separation in the adult and aged group under EE but not the HC condition (**Fig. 1B**). Of the 6-weeks old mice groups, 11 metabolites had a significant difference (*P* < 0.05, unpaired *t*-test) between EE and HC conditions. In contrast to the changes in metabolite levels in adult mice, the changes were more pronounced in aged mice during EE (**Fig. 1C**). As shown in the hierarchical clustering heatmap (**Fig. 1D**), we discovered activity-dependent changes in metabolites in the aged hippocampus that are involved in multiple pathways including glycolysis, pyruvate metabolism, galactose metabolism, TCA cycle, and oxidative phosphorylation.

**Figure 1.**
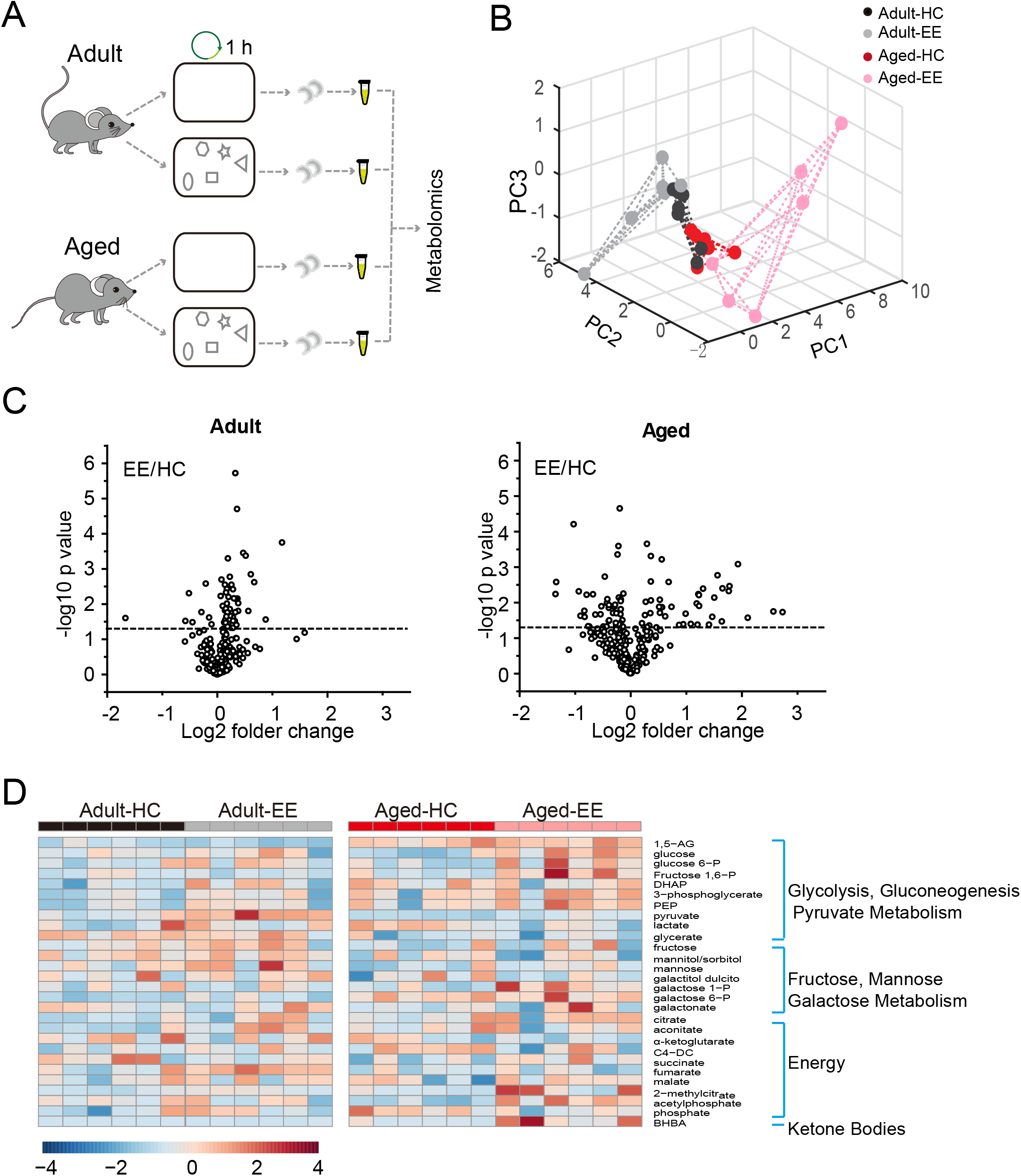
Metabolic analysis of the hippocampus during content exploration in aged mice. A, Experimental design schematic for the metabolic analysis of aged hippocampus. B, PCA analysis of six repeats of adult and aged group underlying home cage (HC) and enriched environment (EE) conditions. C, Volcano plot of the fold-change and p-value for all 54 metabolites between HC and EE conditions. HC, n = 6; EE, n = 6; *P < 0.05; unpaired t-test. D, Hierarchical clustering heatmap of metabolites with significant differences between HC and EE conditions.

Taken together, our metabolomics analysis of the adult and aged hippocampus revealed substantial changes in levels of energy production-related metabolites in the aged mice when compared to those in adult mice during enriched contextual exploration. This analysis suggests a possible energy metabolism shift in the aged hippocampus during a hippocampus-engaged exploratory behavior.

### Decreased glycolytic dynamics in the aged hippocampus during an enriched contextual exploration

Substantial changes in metabolites in the activated hippocampus of both adult and aged groups during enriched contextual exploration motivated us to explore the metabolism of glucose, an essential carbohydrate metabolite. Importantly, the mammalian brain heavily depends on glucose as its main source of energy, although it can adapt to utilize ketone bodies during periods of starvation (Hasselbalch et al., 1994; Mergenthaler et al., 2013). It has been generally considered that the brain glucose metabolism starts with glycolysis in the astrocytes (Belanger et al., 2011). The end product of glycolysis, pyruvate, is then catalyzed into lactate, which is then transported into neurons to enter the TCA cycle for further metabolization. TCA cycle metabolites are then used to generate ATP via the electron transport chain in the mitochondria.

Therefore, we first analyzed the metabolites involved in glycolysis as summarized in **Figure 2A**. The concentrations of glucose and glycolytic metabolites between HC and EE were compared. We found that in the adult hippocampus there was a slight increase of glucose and metabolites involved in glycolysis, including glucose-6-phosphate, DHAP, 3-phosphoglycerate, and phosphoenolpyruvate during enriched contextual exploration (**Fig. 2B-G**). A significant increase of pyruvate was observed in these mice under EE. In contrast, in the aged hippocampus, a substantial increase in hippocampal glucose concentration was observed during contextual exploration (**Fig. 2H**). In turn, levels of glucose 6-phosphate and fructose 1,6-bisphosphate had increased substantially (**Fig. 2C and D**). Interestingly, we discovered that levels of DHAP and 3-phosphoglycerate, which are downstream of glucose 6-phosphate and fructose 1,6-bisphosphate, surprisingly remained unchanged (**Fig. 2E and F**). Furthermore, levels of phosphoenolpyruvate also remained unchanged during EE conditions (**Fig. 2G**). We should point out that an increase in pyruvate (**Fig. 2H**), the end product of glycolysis, was discovered. This suggests a compensation mechanism from other energy sources to supplement the shortage in pyruvate production during glycolysis.

**Figure 2.**
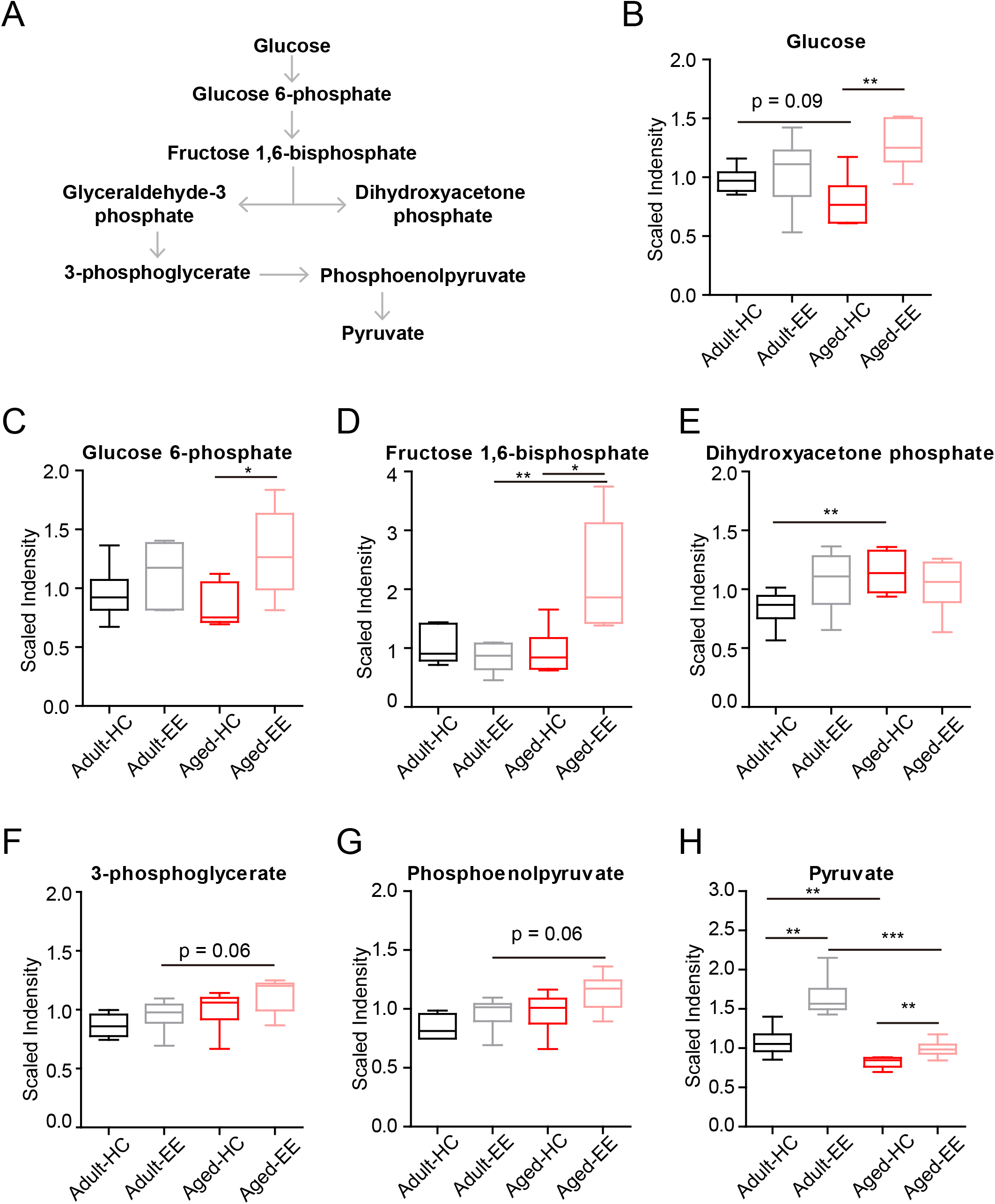
Concentration of metabolites related with glycolysis of aged hippocampus. A, Schematic drawing of the general glycolytic metabolism pathway. B-H, Box plots of the levels of glucose (B), glucose-6-phosphate (C), fructose 1, 6-bisphosphate (D), dihydroxyacetone phosphate (DHAP) (E), 3-phosphoglycerate (F), phosphoenolpyruvate (G) and pyruvate (H) in adult and aged mice under HC and EE condition. HC, n = 6; EE, n = 6; *P < 0.05; unpaired t-test.

Altogether, the analyses revealed that glycolysis, the first step of glucose metabolism for energy production, changed during EE conditions and ended with elevated pyruvate levels in adult mice. However, although the directional dynamics remained similar in the early steps of glycolysis in the aged mice, there was a significant break from the trend in the cleavage from fructose 1,6-bisphosphate to DHAP, suggesting a possible defect in the cellular mechanisms of the aged animals.

### Disrupted cellular respiration pathway in the aged hippocampus in comparison to that of the adult during an enriched contextual exploration

To further analyze the changes in the metabolic pathway between adult and aged mice during enriched contextual exploration, we analyzed the pathway following the production of pyruvate, the end product of glycolysis. Pyruvate serves as an essential substrate for ATP production via the TCA cycle. In the presence of oxygen, the mitochondrion facilitates aerobic respiration via the TCA cycle. Acetyl-CoA is first produced from pyruvate, which then enters the TCA cycle and is then oxidized to CO2 while reducing NAD to NADH and FAD to FADH2, as illustrated in **Figure 3**. NADH and FADH2 are then used by the electron transport chain to generate ATP as part of oxidative phosphorylation. In the TCA cycle, the metabolites of citrate, aconitate, alpha-ketoglutarate, succinate, succinylcarnitine, fumarate, and malate were detected and analyzed in the hippocampus of the four experimental conditions.

**Figure 3.**
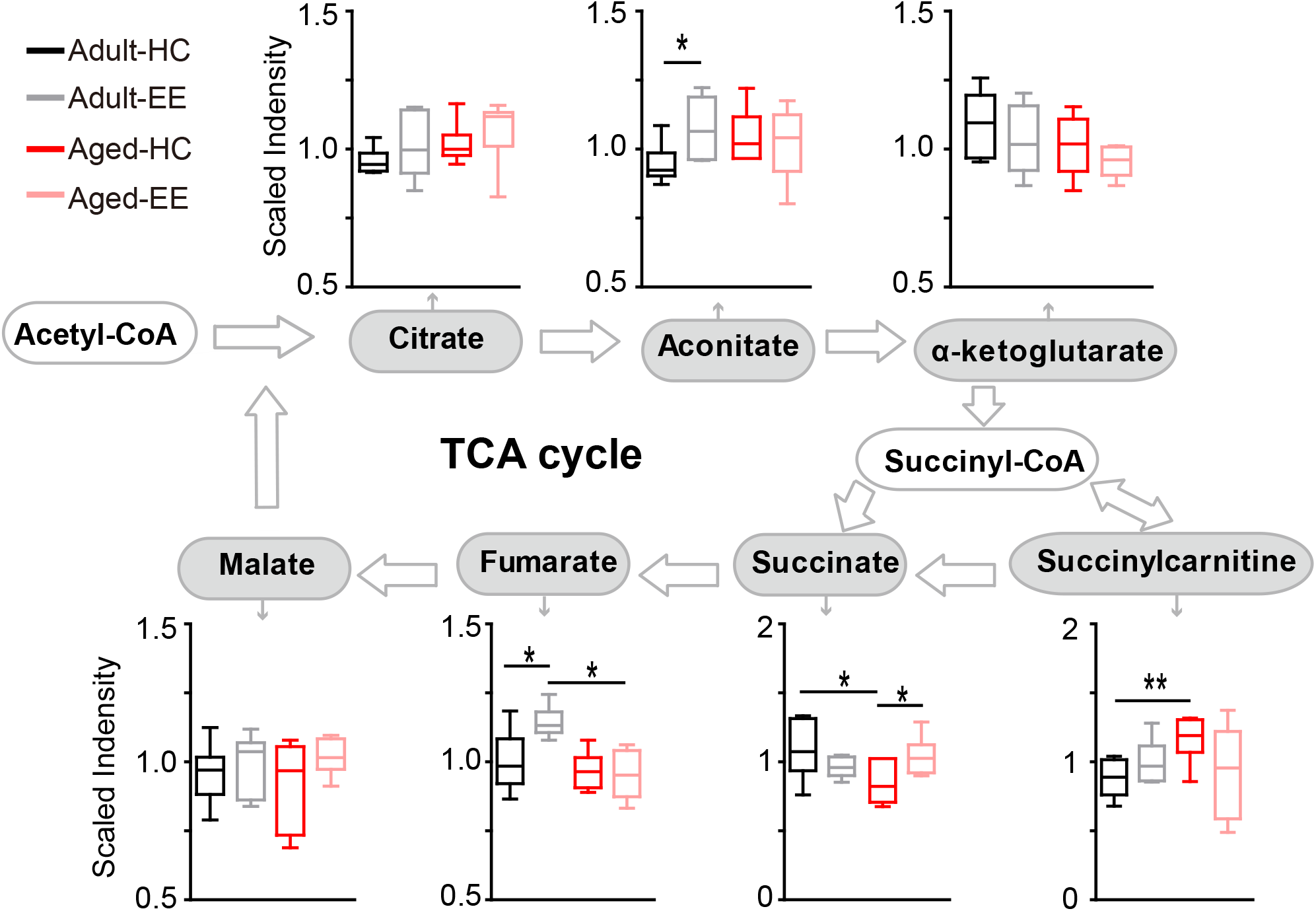
Concentration of metabolites related to the TCA cycle of the aged hippocampus. Box plots of metabolites involved in the TCA cycle show the concentration change between adult and aged hippocampus under HC and EE condition. TCA cycle key metabolites, including citrate, aconitate, alpha-ketoglutarate, succinate, fumarate and malate were plot according to the metabolic pathway.

In the adult hippocampus, a slight increase in metabolite levels involved in the TCA cycle was observed, including aconitate and fumarate during enriched contextual exploration (**Fig. 3**). In contrast, in the aged hippocampus, a visible increase in succinate was found during contextual exploration. Succinyl-CoA was preferentially diverted into succinylcarnitine as opposed to succinate in the aged home cage condition animals as compared that of the adult. This suggests that the TCA cycle does not run as efficiently in aged animals in comparison to adult animals. Succinylcarnitine, a metabolic “dead end”, accumulates in aged control hippocampi, suggesting less energy production via the TCA in aged animals. Importantly, this diverted effect seemed to be “rescued” in aged animals by placing them under EE conditions, leading to increased succinate and decreased succinylcarnitine.

Altogether, the analyses revealed that certain metabolite levels in the TCA cycle significantly increased in adult mice during contextual exploration. However, these significant increases were not paralleled in the aged mice placed under the same conditions. In the aged hippocampus, the TCA cycle may be diverting metabolites to succinylcarnitine, making it less efficient at running aerobic metabolism, suggesting possible additional mechanisms to compensate the short of TCA-based energy production.

### Elevated ketone metabolism in the aged hippocampus in comparison to that of the adult during enriched contextual exploration

In the aged hippocampus, both glycolysis and the TCA cycle seemingly become less efficient during contextual exploration and are unable to supply the energy demands of the brain. The brain might supplement its energy requirements by utilizing ketone bodies when glucose levels are insufficient. Therefore, we set out to measure ketone concentrations in the hippocampus across aging and hippocampal stimulation.

Ketone bodies include acetoacetate and beta-hydroxybutyrate (BHBA). Acetoacetate can be transformed into acetyl-CoA to enter the TCA cycle in the absence of sufficient ATP production from glycolysis and the TCA cycle (**Fig. 4A**). In our metabolomics assay, we successfully detected BHBA (**Fig. 1**). In the adult brain, there were few changes observed in the concentration of BHBA during contextual exploration. However, a substantial increase in BHBA concentration was observed in the aged brain during contextual exploration (**Fig. 4B**). The oxidation of BHBA eventually leads to the production of acetyl-CoA, which requires the transfer of CoA from succinyl-CoA. This may result in a higher concentration of succinate (**Fig.3**). This is a possible cause of the increase in succinate observed in the TCA cycle.

**Figure 4.**
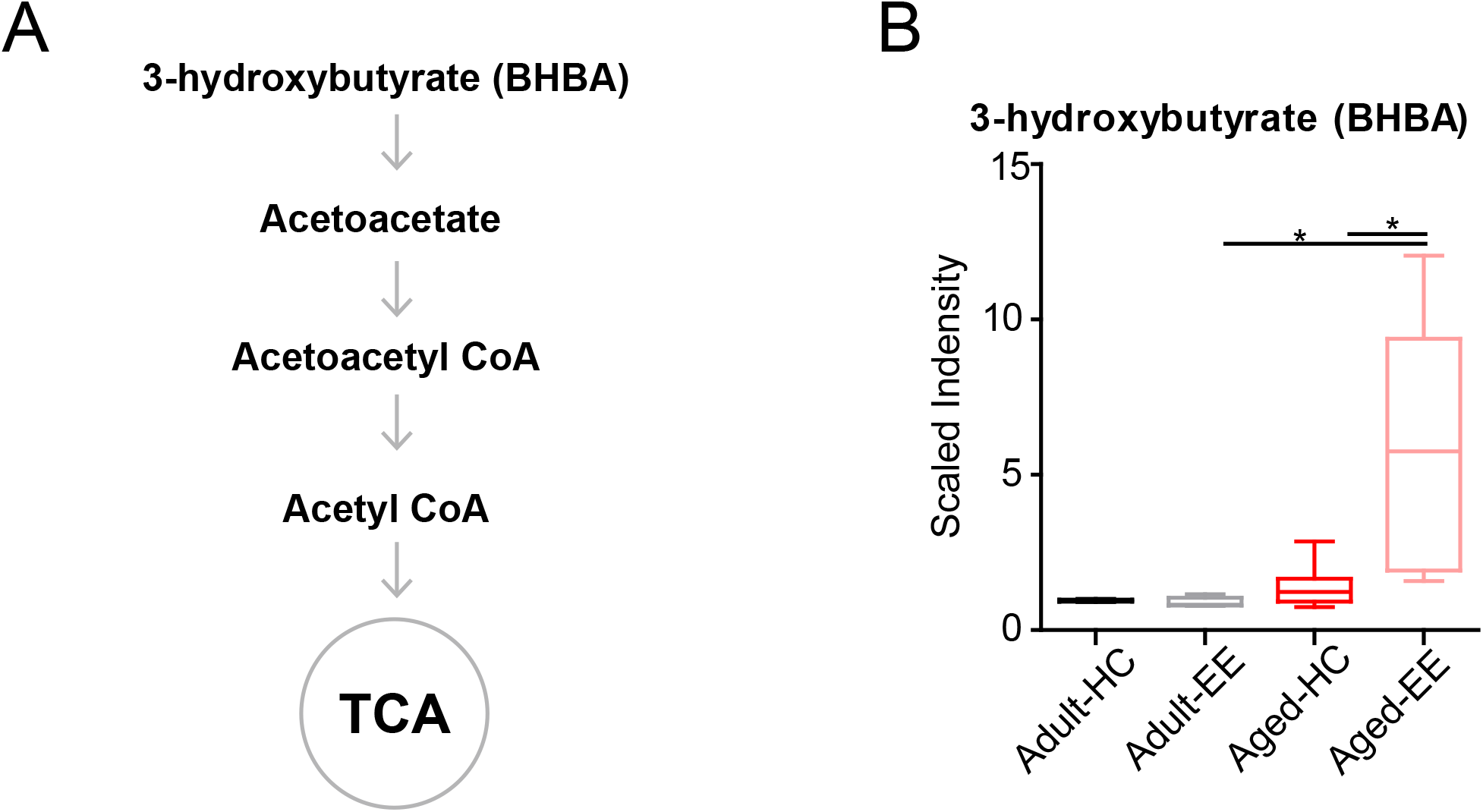
Concentration of metabolites related to acetyl CoA of aged hippocampus. A, Schematic drawing of metabolic pathway including ketone body metabolism linking with acetyl CoA. B, Box plot of the levels of 3-hydroxybutyrate (BHBA) in adult and aged mice. HC, n = 6; EE, n = 6; *P < 0.05; unpaired t-test.

Altogether, the analyses revealed that the level of the ketone body BHBA remains comparable in the aged brain as compared to the adult brain during HC. Additionally, there was no change in the level of BHBA in the adult brain between HC and EE. However, the level of BHBA in the aged hippocampus is substantially elevated during EE.

## Discussion

How does energy metabolism change in the aged brain, especially during cognitive activity? In this study, using a metabolome analysis method, we measured the changes in the levels of various metabolites under enriched contextual exploration conditions in both the adult and aged hippocampus. The entire metabolite profiling displayed substantial changes in concentrations of metabolites involved in energy production pathways. We first explored changes in glucose metabolism by analyzing the levels of metabolites involved in the glycolytic process. The levels of certain metabolites experienced significant changes and we discovered an overall decrease in efficiency of pyruvate production in the aged brain. Furthermore, we analyzed levels of metabolites in the TCA cycle, finding an increase in succinate in the aged brain. This suggests a secondary pathway, namely from ketone bodies, that bolstered the concentration of succinate. Together, our results suggest impairments in specific enzymes of metabolic pathways that lead to the overall decrease in energy metabolism in the aged brain at baseline and during contextual exploration.

During hippocampus-engaged behaviors, it has been shown that glucose levels decrease substantially in the extracellular compartment (McNay et al., 2000; McNay et al., 2001), and glucose utilization is increased to meet the elevated neural activities (De Bundel et al., 2009; Belanger et al., 2011; Figley and Stroman, 2011), suggesting elevated glucose-related metabolism. Thus, to better investigate this, we analyzed the levels of various metabolites in the glucose-related metabolism to gauge the changes in these levels during hippocampus-engaged behaviors. Using metabolomic analyses, we found that EE led to changes in the concentration of a variety of energy production-related metabolites in the adult hippocampus. Interestingly, in the aged hippocampus, we observed an apparent diversion of pyruvate and acetyl-CoA toward ketone metabolism in the aged brain, especially under conditions of enriched environmental exploration, as compared to the adult brain.

During glycolysis, although there was a substantial increase in pyruvate concentration during the EE condition in the aged hippocampus, the absolute levels of pyruvate between the adult and aged hippocampus show a lower metabolism efficiency in the aged hippocampus (**Fig. 2H**). While there were no significant differences in the levels of glucose between the adult and aged hippocampus under both HC and EE conditions, there were major differences in the levels of pyruvate under both conditions. In the aged hippocampus, the glucose concentration was substantially increased under EE, possibly reflecting increased blood glucose delivery to the hippocampus, but impairment in the ability to utilize this glucose for aerobic metabolism. Previous studies have shown that the aged hippocampus increases glycogen metabolism enzyme concentrations and shifts in these enzymes’ localization from astrocytes to neurons (Drulis-Fajdasz et al., 2018). However, given the decrease of pyruvate in the aged cohort despite a substantial increase in glucose levels, we hypothesize a decrease in efficiency of glucose metabolism in the aged brain. Future work should aim to specifically measure glucose consumption in different brain areas across ages, for example, by measuring glucose transporter kinetics and glucose labeling to trace the accumulation of metabolic byproducts.

In the TCA cycle of the adult brain, there was an increase in the levels of aconitate and fumarate during the EE condition. In the TCA cycle of the aged brain, there was a substantial increase in succinate concentration during EE conditions (**Fig. 3**). The question remains regarding the cause of this buildup of succinate. One possibility is that succinate dehydrogenase is less efficient in the aged brain as compared to the adult brain. Alternatively, we hypothesize that this increased succinate may result as a byproduct of BHBA metabolism. In our further analysis, we found that BHBA levels had a sharp, significant increase from the aged HC to the aged EE condition (**Fig. 4B**). Since the breakdown of BHBA to acetyl-CoA involves changing succinyl-CoA to succinate, this may serve as a possible mechanism to explain the significant increase in succinate in the aged cohort.

Neurodegeneration is the progressive decline of functionality due, in part, to alterations in cellular metabolism and respiration (Procaccini et al., 2016). The functionality of the brain fundamentally relies on energy that is metabolized from various sources. This current study adds to a growing body of literature using a metabolomics approach to understand the energy supply dynamics. This work provides insights into metabolic derangements associated with aging and degeneration which may lead to the development of therapeutic strategies to help mitigate such degeneration.

